# Knee Joint Biomechanics During Lunges at Different Tibial Angles and External Loads: A Musculoskeletal Analysis with Finite Element Insights

**DOI:** 10.64898/2026.08.07.743401

**Authors:** Lidong Gao, Shunxiang Gao, Zhenghui Lu, Gusztáv Fekete, Zixiang Gao

**Affiliations:** Multidisciplinary Doctoral School of Engineering Sciences, Széchenyi István University, 9026 Gy ̋or, Hungary; Faculty of Sports Science, Ningbo University, Ningbo, China; Research Academy of Grand Health, Ningbo University, Ningbo, China; Faculty of Engineering, University of Pannonia, Veszprém, Hungary; Faculty of Kinesiology, University of Calgary, Calgary, Canada

**Keywords:** Lunge, Knee joint, Musculoskeletal model, Finite element analysis, Biomechanics

## Abstract

**Objective:** This study investigates knee joint biomechanics during lunges under varying tibial angles and external loads using musculoskeletal modeling and finite element analysis. The goal is to provide a biomechanical basis for understanding knee loading patterns and optimizing sports training and rehabilitation.

**Methods:** Twenty-six healthy young men performed lunges under tibial inclination angles relative to the ground (60° and 90°) and two external load conditions (bodyweight and an additional 98 N external load). Kinematic and kinetic data were captured using motion capture and force plates. Musculoskeletal models were used to estimate joint moments, range of motion, and stiffness, with data analyzed using two-way repeated-measures ANOVA. Finite element analysis was performed at 90° tibial angle to evaluate tissue stress and displacement.

**Results:** The joint moment at a 60° tibial angle was much higher than at a 90°. External load showed significant effects on knee stiffness, with lower rotational stiffness in the horizontal plane (P < 0.001) and lower coronal plane stiffness at 90° (P = 0.012) under the 98 N external-load condition, indicating reduced resistance to angular displacement in these planes. Under the 90° tibial-angle condition with external load, peak stress and displacement were concentrated in the posterior horn of the meniscus, with a maximum displacement of 3.12 mm.

**Conclusion:** The anterior tilt of the tibia increased sagittal-plane knee loading, while external load mainly reduced joint stiffness in the coronal and horizontal planes. Under the 90° loaded condition, the concentration of stress and displacement in the posterior horn of the meniscus suggests a mechanically unfavorable loading pattern rather than direct evidence of injury risk. These findings may provide useful biomechanical information for load management during lunge-based training and rehabilitation.

## 1. Introduction

The knee joint is one of the most complex and heavily loaded joints in the human body, primarily consisting of the tibiofemoral and patellofemoral joints. Its complicated internal structures, such as bones, menisci, cartilage, and ligaments, are essential for maintaining joint function and three-dimensional stability under both static and dynamic pressures (Felson 2013). As a representative functional exercise, the lunge not only effectively strengthens lower-limb muscles but also plays a critical role in early rehabilitation following anterior cruciate ligament (ACL) reconstruction and fall-prevention training for the elderly (Alkjær et al. 2020; Escamilla et al. 2008; Riemann et al. 2012; Wu et al. 2020). Recent evidence also supports the use of progressive strength-training program to improve muscular strength, endurance, physical self-efficacy, and physical activity participation in middle-aged and older adults (Ward et al. 2026). Therefore, understanding knee joint loading during lunges is important for both athletic training and clinical rehabilitation.

During a lunge, the knee joint is exposed to substantial external loads that generate large joint moments, while coordinating muscle groups to maintain stability (Riemann et al. 2012). This is particularly true under high-load conditions or increased lunge depth, where the knee bears a greater mechanical demand. Knee pain is common in physically active populations, particularly in activities involving repetitive loading of the lower limbs. Recent injury surveillance in amateur football further highlights the relevance of knee loading in physically active populations (Arias et al. 2025). Among these conditions, patellofemoral pain syndrome (PFPS) is one of the most frequently reported forms of anterior knee pain, with a reported annual prevalence of over 20% in the general population and physically active individuals (Smith et al. 2018; van Leeuwen et al. 2023). Clinical studies have demonstrated that improper movement technique or poor load management can generate abnormal joint forces, inducing soft tissue injuries such as patellofemoral pain syndrome (Chuter and de Jonge 2012; Escamilla et al. 2008). Furthermore, long-term repetitive exposure to deleterious mechanical loads may disrupt tissue mechanical homeostasis and eventually lead to cartilage degeneration (Andriacchi et al. 2004).

Previous studies have used motion capture systems, force plates, and musculoskeletal modeling to analyze the macroscopic biomechanical features of the knee during lunges (Escamilla et al. 2008; Gao et al. 2022; Riemann et al. 2012). These approaches can estimate joint kinematics, joint moments, muscle forces, and joint stiffness, which are useful for evaluating movement strategy and mechanical stability (Mei et al. 2022; Rajagopal et al. 2016). In particular, joint stiffness provides an indirect indicator of the knee joint’s ability to resist angular displacement under different loading conditions. However, direct measurement of internal stress distribution in vivo remains difficult due to technical limitations. Traditional musculoskeletal modeling can estimate joint-level mechanical outputs; however, it cannot reveal localized stress and displacement within specific knee tissues, such as the cartilage and menisci.

In contrast, finite element analysis (FEA) allows for the simulation of tissue-level mechanical responses in complex anatomical structures. FEA provides intuitive graphical results that clarify the distribution and magnitude of local loads caused by changes in posture and biomechanics (Yan et al. 2024). Therefore, integrating musculoskeletal modeling with FEA provides a multiscale framework for linking whole-joint loading patterns with local tissue stress and displacement responses.

The primary objective of this study was to investigate the effects of tibial angle and external loading on knee joint biomechanics during lunges using an integrated musculoskeletal–FEA framework. Specifically, musculoskeletal modeling evaluated joint ROM, moments, and stiffness across the four tibial-angle and loading conditions, while FEA examined tissue stress and displacement under the 90° tibial-angle condition. This study proposes two hypotheses: (1) Tibial angle and external loading would significantly affect knee joint moments and stiffness. (2) Under the 90° loaded condition, the external load will greatly increase the stress levels in the knee tissues. Stress and deformation will become more concentrated in certain regions, like the posterior horn of the meniscus. The findings may provide a biomechanical basis for optimizing lunge-based training and rehabilitation strategies.

## 2. Methods

### 2.1 Participants

We used G*Power software (Version 3.1.9.4) to figure out the sample size. A total of 26 individuals were recruited, with a statistical power of 0.80, a significance level of 0.05, and an effect size of 0.40, which was based on similar experimental designs and previous investigations (Faul et al. 2007; Nakagawa and Cuthill 2007). **Table 1** shows all of their specific traits. To ensure data consistency, participants followed a strict pre-test protocol: no food intake two hours before the experiment, and no alcohol or caffeine consumption within 24h. Additionally, participants were required to abstain from any form of physical exercise for 48h prior to testing. All recruited subjects were free of lower limb diseases or injuries for at least six months before the study. The dominant leg was defined as the preferred limb used for kicking a ball. During testing, all participants wore the same style of shoes and compression clothing to minimize any influence from footwear or apparel. The participant chosen for the FE modeling (age: 27 years; height: 1.74 m; weight: 70 kg) was a typical member of this group. The finite element model was based on this representative individual whose characteristics were similar to the group average (Kiapour et al. 2014a); therefore, the FE analysis should be viewed as a case study providing mechanistic insights within the broader group findings, rather than a population-wide estimate. Everyone who took part in the study knew exactly what it was for and how the tests would be done. Before the experiment, each participant gave their written agreement. This study was performed with complete ethical approval and in compliance with informed consent protocols. (TY2025087)

**Table 1.** Basic information of participants.

|  |  |
| --- | --- |
| Age (years) | 26.50±1.50 |
| Height (m) | 1.75±0.05 |
| Weight (kg) | 71.25±7.58 |
| Leg length (m) | 0.89±0.06 |
| Shoulder width (m) | 0.34±0.02 |

### 2.2 Instruments

Kinematic data were captured using a Vicon motion capture system (Oxford Metrics, Ltd., Oxford, UK) equipped with 10 high-speed infrared cameras and Nexus analysis software, operating at a sampling frequency of 200 Hz. Two 3D force plates (Kistler, Winterthur, Switzerland) recorded kinetic data at the same time at a rate of 2000 Hz. To assess the plausibility of muscle activation patterns estimated by OpenSim static optimization, electromyography (EMG) signals from the rectus femoris, biceps femoris long head, tibialis anterior, and lateral gastrocnemius were recorded using a Delsys EMG system (Delsys, Boston, MA, USA) at a frequency of 1000 Hz. The motion capture and force-plate data were hardware-synchronized through the Vicon Nexus acquisition system. Because the EMG system could not be directly synchronized with the Vicon-force plate system, EMG recordings were temporally aligned during post-processing using an external video reference and common lunge events.

Based on CT and MRI data from one representative participant, a finite element model of the knee was constructed. The CT scans were performed using a Somatom Emotion 16-slice spiral CT scanner, with a reconstruction layer thickness of 1 mm and a spacing of 0.5 mm (settings: 120 kV, 250–300 mA). For internal soft tissue visualization, MRI scans were conducted using a 1.5T high-field MRI system (United Imaging Healthcare), equipped with a built-in body coil. The MRI data were acquired in the sagittal plane with a layer thickness of 1mm and no inter-slice gaps.

### 2.3 Procedures

Before the tests, a standard tape measure was used to measure the leg length of every individual. The distance from the anterior superior iliac spine to the medial tibial condyle was used to measure leg length. The length of the lunge step was set at 70% of every individual’s leg length (Gao et al. 2022). Tape markings were put on the beginning point and the target force plate to make sure that all trials were the same. To make the stance width the same for everyone, the participant’s shoulder width was measured and marked on the floor and force plates as the distance between the feet. Additionally, we recorded the time taken for each participant to perform a single lunge. The average time for all participants was then set as the standard time for all experimental trials to make sure that everyone moved at the same speed.

Before the testing began, participants performed a 5-minute warm-up on a stationary treadmill (h/p/cosmos sports and medical GmbH, Nussdorf-Traunstein, Germany) to ensure they were in an active physical state. Under the guidance of professional staff, each participant practiced the lunge movements for all experimental conditions. Verbal instructions were provided to ensure they were fully familiar with the movement sequence and could perform it safely and consistently. The position of the lower leg was chosen as a control variable to make distinct lunge depths. The tibial angle, which is the angle between the tibia’s long axis and the ground, was used as the main independent variable to make the lunge depth the same. It was set at 60° for a deeper lunge and 90° for a shallow lunge.

During the familiarization trials, a rope positioned at the target angle was used to help participants practice and achieve the required tibial position. During data collection, the target angles were monitored using floor reference marks, a side-positioned goniometer, and continuous visual feedback from an investigator. The achieved tibial angle was further verified post hoc using motion-capture data by calculating the orientation of the tibial long axis relative to the laboratory ground plane at the target lunge position. Trials were accepted only when the measured tibial angle fell within ± 5° of the target angles. The achieved tibial angles were 58.6 ± 3.4° for 60° condition and 87.5 ± 2.1° for the 90°. During the external loads test, each participant held a dumbbell with a mass of 5 kg in each hand, corresponding to a total additional mass of 10 kg and an external gravitational load of approximately 98 N.

Before the EMG sensors were put on, the skin was prepared by shaving off any hair and cleaning off oils, sweat, and other residue. This was done to make sure that the signal was of high quality. Electrode placement followed the SENIAM recommendations where applicable (Hermens et al. 2000). Based on tangible bony features, these sensors were placed over each target muscle’s muscle belly with electrodes parallel to muscle fibers. The electrodes for the long head of the biceps femoris (BF) were placed in the middle of the muscle between the lateral femoral epicondyle and the ischial tuberosity. The electrodes for the rectus femoris (RF) were placed in the middle of the line that connects the anterior superior iliac spine to the superior border of the patella. The sensors for the tibialis anterior (TA) were inserted at the proximal one-third of the line between the fibular head and the medial malleolus. The sensors for the lateral gastrocnemius (LG) were positioned at the proximal one-third of the line between the fibular head and the calcaneus. Before the lunge trials, maximum voluntary isometric contraction (MVIC) tests were performed for the rectus femoris, biceps femoris long head, tibialis anterior, and lateral gastrocnemius using a dynamometer (CON-TREX MJ System, CMV, Dübendorf, Switzerland). Each participant performed three 5s MVIC trials for each muscle, with a 1 min rest interval between trials. The highest EMG amplitude obtained during the MVIC trials was used as the reference value for EMG normalization. **Figure 1** shows the integrated framework for motion capture-based musculoskeletal and FE modeling.

**Figure 1.**
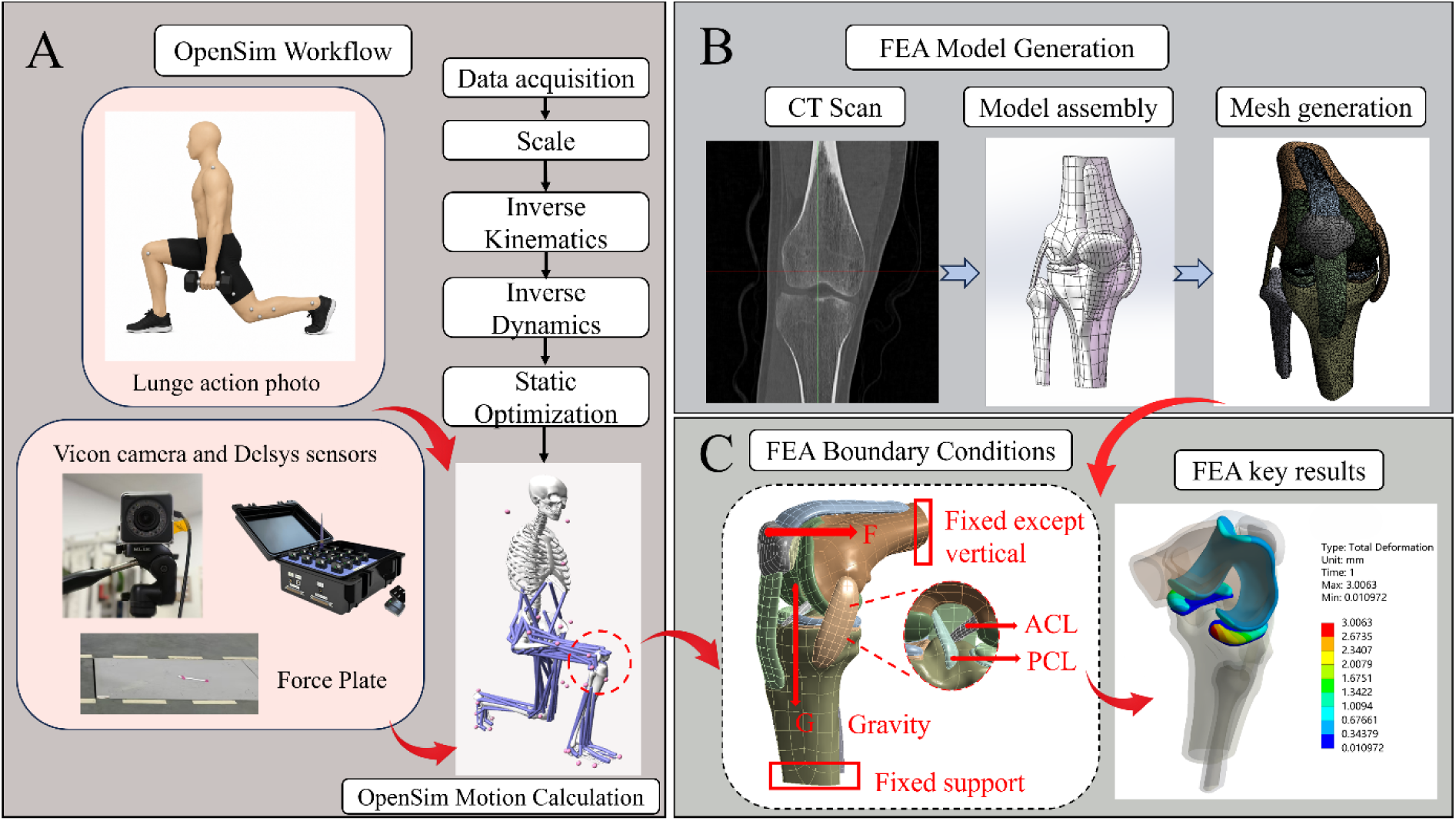
A: Motion capture data acquisition and the OpenSim workflow; B: The FE modelling process. C: Diagram illustrating the details about anterior cruciate ligament (ACL) and posterior cruciate ligament (PCL) and load application conditions and results in FE model.

Each participant performed four lunge conditions: bodyweight at 60° and 90°, and external load (10 kg) at 60° and 90°. The external load conditions and lunge angles were thoroughly randomized for each participant, and right leg data were obtained across all four conditions. The final dataset includes six successful trials for each condition. A five minutes rest time followed each lunge condition to avoid muscular fatigue from impacting results.

### 2.4 Data collection and processing

The motion capture data acquisition and OpenSim musculoskeletal modeling workflow are illustrated in **Figure 1A**. Each lunge phase was the time between the frame immediately preceding right foot contact with the force plate and the participant’s return to standing. For EMG alignment, the external video reference and common movement events, including right-foot contact and the return-to-standing phase, were used to temporally match the EMG recordings with the Vicon-force plate data as closely as possible. Sagittal, coronal, and horizontal kinematic and kinetic data were gathered to investigate knee biomechanical changes under different conditions. A trial was successful if the right heel made initial contact with the ground on the marker line, the trailing leg knee approached the ground without contact, the trunk remained as upright as possible, and the right knee reached the target angle without significantly exceeding it. The dataset excludes trials where the right toe or trailing knee hit the ground.

Initial processing of the captured data was performed using Vicon Nexus. The marker trajectories and ground reaction forces were filtered using a zero-lag, fourth-order Butterworth low-pass filter with cutoff frequencies of 12 Hz and 30 Hz, respectively. A threshold of 20 N was applied to the ground reaction force data, and any values below this limit were excluded. Using Matlab R2018a (The MathWorks Inc., Natick, MA, USA), the C3D files were converted into formats compatible with OpenSim 4.3 (.mot and .trc) and subsequently imported into the OpenSim software for further data processing (Zhou et al. 2021).

The model used in this study was adapted from the original OpenSim model (Rajagopal et al. 2016) as modified by Mei et al. (Mei et al. 2022), a version previously validated in several studies (Gao et al. 2023; Mei et al. 2019; Yu et al. 2023). During static calibration, the model was scaled based on each participant’s marker positions and body weight. We ensured model accuracy by adjusting the marker positions until the root mean square (RMS) error between experimental and virtual markers was below 0.02. Kinematic data were then calculated using the Inverse Kinematics (IK) tool, while net knee joint moments were determined via the Inverse Dynamics (ID) algorithm. The ID tool solves the classical equations of motion to derive the resultant forces and torques at each joint.

Normalizing peak joint moment data to 101 points ensured graphical consistency. All joint moments were standardized by body weight to account for individual variances and remove body weight (Hsue et al. 2009). Each condition’s joint stiffness was determined by calculating the joint moment range (ΔM) using maximum and minimum values. This formula estimated stiffness:

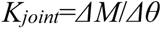

*K_joint_* represents joint stiffness (units: Nm/kg/deg), *ΔM* is the variance of joint moment, and *Δθ* is the range of joint motion.

Individual muscle forces were estimated using Static Optimization (SO). The SO tool solves equations of motion for unknown generalized forces like joint torques using the model’s known motion as an extension of inverse dynamics. Net joint moments are further decomposed into muscle forces at each instantaneous time point. To assess the plausibility of the SO-derived muscle activation patterns, the experimentally recorded EMG signals were compared with the OpenSim-estimated muscle activations. Raw EMG signals were processed offline using a 10-500 Hz fourth-order Butterworth band-pass filter, full-wave rectification, and a 50 ms moving root mean square (RMS) window. The processed EMG amplitudes during the lunge trials were normalized to the corresponding peak MVIC value for each muscle and expressed as %MVIC. Both the normalized EMG envelopes and SO-derived muscle activations were time-normalized to 101 points across the lunge phase for waveform comparison. Waveform agreement was then quantified using the maximum cross-correlation coefficient and normalized root mean square error (nRMSE).

### 2.5 Development of a FE model

The knee joint model was reconstructed to obtain the geometric structures of the bones, cartilage, and ligaments. The MRI data were utilized to distinguish the anatomical boundaries for the articular cartilage (femoral, tibial, and patellar), the medial and lateral menisci, and the main ligaments, such as the ACL, posterior cruciate ligament (PCL), lateral collateral ligament (LCL), medial collateral ligament (MCL), and patellar tendon (PT). Furthermore, the specific bone attachment points for each major ligament were identified (Al Mohammad and Gharaibeh 2024; Innocenti et al. 2016). The three-dimensional reconstruction was performed using MIMICS 21.0 (Materialise, Leuven, Belgium). The resulting models were refined and smoothed in Geomagic Studio 2021 (Geomagic, Inc., Research Triangle Park, NC, USA) before being assembled in SolidWorks 21 (SolidWorks Corporation, MA, USA).

The final mesh generation was performed using Ansys Workbench 2021 R1 (ANSYS Inc., Canonsburg, PA, USA). The bones, cartilage, menisci, and ligaments were discretized using 3D continuous solid linear tetrahedral elements. To balance computational efficiency with geometric precision, the element sizes were set to 3mm for the bones, 1mm for the menisci and cartilage, and 1.5mm for the ligaments (Beidokhti et al. 2017; Gao et al. 2025). To ensure the model results remained insensitive to mesh density, we applied mesh refinement in areas with high stress gradients to improve local accuracy. By comparing the trends of stress, strain, and contact variables across different mesh densities, we found that the convergence differences for the selected mesh were within an acceptable range (less than 5%), demonstrating an optimal balance between accuracy and computational efficiency (Kiapour et al. 2014a; Li et al. 1999; Rooks et al. 2022).

To obtain reliable mechanical data while maintaining computational efficiency, simplified material models were used to describe the mechanical behavior of various knee tissues. This simplification is justified as the elastic modulus of bone much exceeds that of soft tissue; thus, its minor deformations have little impact on the overall stress distribution of the joint (Haut Donahue et al. 2002). For the cartilage, menisci, and ligaments, these tissues were also simplified to make the simulation run faster and more stable by treating them as isotropic and linear elastic materials (Galbusera et al. 2014; Yan et al. 2024). Although cartilage, menisci, and ligaments are viscoelastic and may exhibit strain-rate-dependent behavior, isotropic linear elastic models were used to improve numerical stability and computational efficiency. The final FE model included 402881 nodes and 236677 elements. We assigned material features based on values found in the literature, with **Table 2** (Lu et al. 2025; Lu et al. 2023; Rao et al. 2024; Yan et al. 2024) giving more information about the specific parameters. **Figure 1BC** show the process of establishing the finite element model and the schematic diagram of the operating conditions under which the model is subjected to load.

**Table 2.** Material parameters of the model components.

|  | Young's Modulus<br>(MPa) | Poisson's ratio<br>(v) | Mesh Size<br>(mm) |
| --- | --- | --- | --- |
| Cortical bone | 12000 | 0.30 | 3.0 |
| Cancellous bone | 500 | 0.30 | 3.0 |
| Cartilage | 15 | 0.46 | 1.0 |
| Meniscus | 59 | 0.45 | 1.0 |
| ACL | 116 | 0.30 | 1.0 |
| PCL | 87 | 0.30 | 1.0 |
| MCL | 48 | 0.30 | 1.5 |
| LCL | 48 | 0.30 | 1.5 |
| PT | 87 | 0.30 | 1.0 |

### 2.6 Boundary conditions and loads

The distal ends of the tibia and fibula were fixed to set the boundary conditions. The proximal femur, on the other hand, could only move vertically. Given the high geometric complexity of the knee model and the large number of elements, the contact interfaces were simplified to improve convergence robustness under high-load and large-deformation conditions(Yan et al. 2024). For the menisci, bonded contact was defined between the inferior meniscal surfaces and the tibial cartilage, whereas a no-separation contact formulation was applied between the superior meniscal surfaces and the femoral cartilage. This configuration constrained the menisci on the tibial plateau while allowing compressive load transfer from the femoral cartilage during the simulated lunge posture. This approach was adopted to balance numerical stability with the representation of the primary tibiofemoral load-transfer pathway.

The loads used were exactly the same as the results from the SO tool, with the quadriceps’ force being applied to the top edge of the patella. Gravitational and external loads, ascertained from the exact angles and timestamps in the IK data, were administered at the midpoint of the femoral condyles, adhering to established guidelines from prior research (Beidokhti et al. 2017; Navacchia et al. 2018; Yang et al. 2024). The simulation employed a multi-load-step approach, enabling significant deformation effects to guarantee numerical stability and computational accuracy for soft tissues subjected to high-load contact conditions.

### 2.7 Model Simulation and Validation of FE Models

SolidWorks was used to anatomically assemble the three-dimensional solid knee model in this investigation. Real studies cannot use real-time dynamic imaging of bone orientation, so we used OpenSim IK data for posture alignment. To address the differences in global coordinate system definitions between the OpenSim musculoskeletal environment and the SolidWorks FE environment, a cross-platform coordinate mapping mechanism was established. The IK output flexion-extension, internal-external rotation, and adduction-abduction values were properly translated onto the functional rotation axes by establishing the femur’s local anatomical axes relative to the tibia. Anatomical landmarks were used to modify the patellar position to seat it in the femoral trochlear groove and ensure the patellar tendon (PT) was taut. This method assures that the static FE model’s spatial orientation matches the experimental dynamic postures.

To validate the biomechanical reliability of the FE model, we conducted a simulated anterior drawer test, a clinically recognized validation method. In this procedure, the femur was completely fixed while a standard 134 N anterior load was applied to the tibial plateau. This setup was designed to simulate the response of the ACL, MCL, and LCL under anterior shear forces (Pena et al. 2006; Ren et al. 2022). We then compared the model’s predicted anterior tibial translation with experimental data to verify its validity. The results showed that both the anterior translation of the tibia and the peak von Mises stress distributions for the ACL, MCL, and LCL were highly consistent with previously reported cadaveric data and similar simulation studies (Benos et al. 2020; Ren et al. 2022; Shao et al. 2022).

While the validation shows that the model is biomechanically reliable under baseline external load conditions, in vivo biomechanical research under extreme combination situations is difficult to get experimental data for. Relevant experimental comparison data for the 60° condition employed in this investigation is lacking in the literature. Complex soft tissue self-contact and significant mesh distortion make nonlinear numerical iterations difficult to converge at very high flexion angles. Because of the absence of directly comparable experimental data in the literature, accurate validation under these conditions is challenging. The most common functional weight-bearing angle is 90° tibial angle. This balance of computational dependability with biomechanical realism minimizes computing costs and ensures that the model’s stress distribution characteristics match the load magnitudes in Escamilla et al. (Escamilla et al. 2008). Thus, the model may offer valuable clinical reference for future research and applications.

### 2.8 Statistical Analysis

This study employed IBM SPSS Statistics 26 (SPSS, Chicago, IL, USA) to analyze range of motion (ROM), peak moment, and peak stiffness in the sagittal, coronal, and horizontal planes. In this study, the directions of movement in the three planes are delineated as follows: The sagittal plane shows knee flexion (+) and extension (–), the coronal plane shows adduction (+) and abduction (–), and the horizontal plane shows internal rotation (+) and external rotation (–). The external loads, including quadriceps force and gravitational load, applied to the FE model were derived as the mean values from experimental data. A two-way repeated measures ANOVA was conducted to determine the main effects and interactions of the tibial angle (60° and 90°) and external load condition (body weight and 98N external load) on peak joint moments and stiffness. Prior to the analysis, the normality of the data was confirmed using the Shapiro-Wilk test. If the data were found to deviate from normality, the Wilcoxon signed-rank test was employed for pairwise comparisons. Since each factor had only two levels, the assumption of sphericity was inherently satisfied. Post-hoc analysis was performed using Bonferroni-corrected pairwise comparisons. The results are presented as Mean ± SD, and statistical significance was set at P < 0.05.

## 3. Results

### 3.1 Model evaluation

As illustrated in **Figures 2A–D**, the MVIC-normalized EMG envelopes were compared with the OpenSim static optimization-derived muscle activation curves. Quantitative waveform comparison was performed using the maximum cross-correlation coefficient and nRMSE. The maximum cross-correlation coefficient and nRMSE were 0.280 and 0.362 for the lateral gastrocnemius, 0.588 and 0.360 for the biceps femoris long head, 0.578 and 0.286 for the rectus femoris, and 0.640 and 0.241 for the tibialis anterior, respectively. These results indicate variable agreement between EMG and SO-derived activations. Therefore, EMG comparison was used only as a supplementary plausibility check of SO-derived muscle activation patterns, rather than as strict model validation or validation of FE-derived stress and displacement. To validate the FE model, a simulated anterior drawer test was performed at 0° of flexion by applying a 134 N anterior force to the tibia (**Figure 2E**).

**Figure 2.**
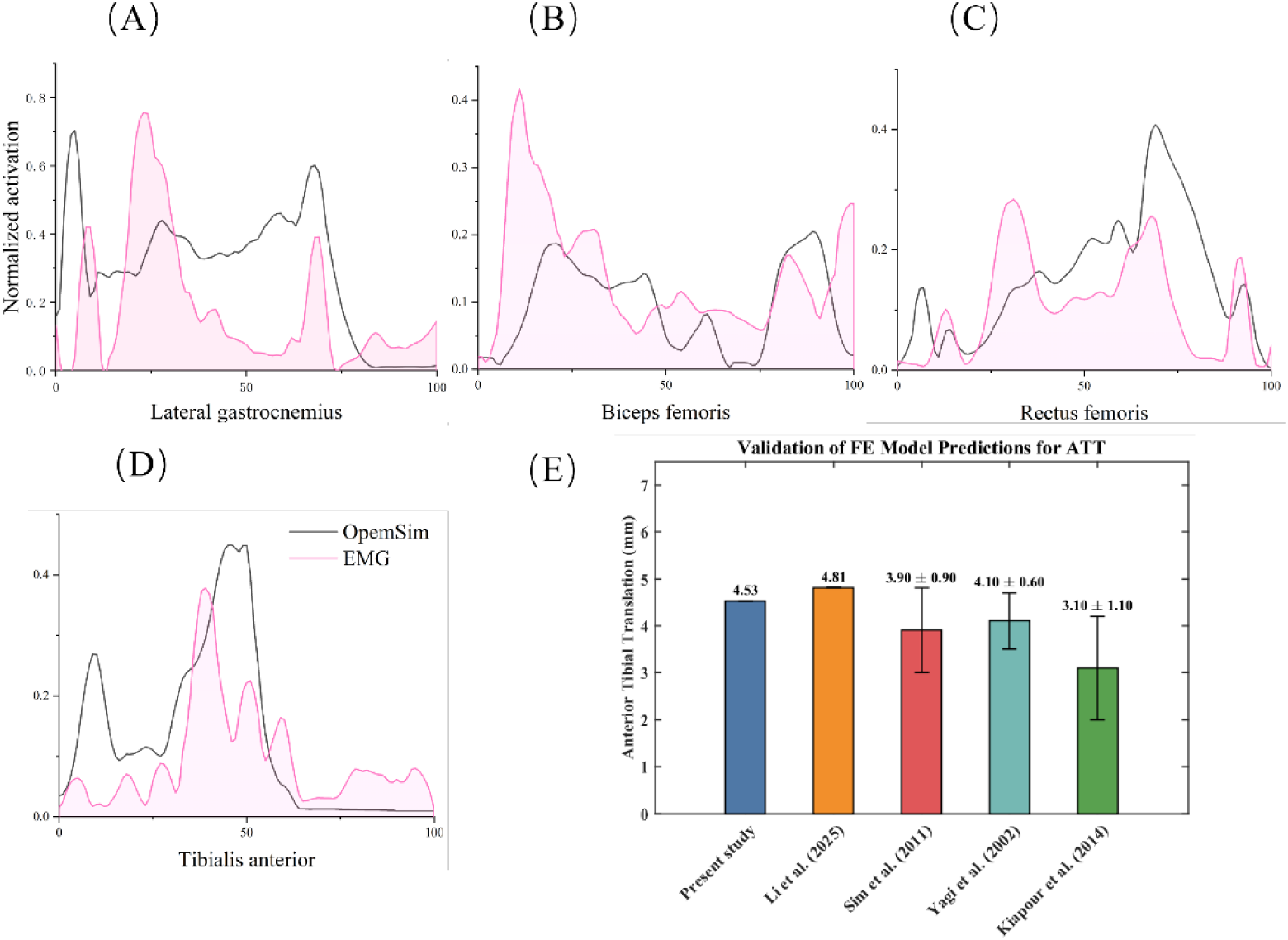
Comparison of MVIC-normalized EMG envelopes and OpenSim static optimization-derived muscle activations for the lateral gastrocnemius (A), biceps femoris (B), rectus femoris (C), and tibialis anterior (D); Validation of the FE model against results from other studies (E).

The model predicted an anterior tibial translation of 4.50 mm, which is within the reasonable range reported in previous studies (Kiapour et al. 2014b; Li et al. 2025; Yagi et al. 2002) and falls under the clinically accepted displacement threshold for a healthy knee (< 6 mm). This result, combined with the previously mentioned load magnitude comparisons, confirms that the coupled framework developed in this study possesses rigorous biomechanical authenticity when handling the high-intensity conditions of a lunge. This moderate compliance more accurately reflects the biomechanical characteristics of living subjects and is beneficial for simulating the large-deformation response of soft tissues at deep flexion angles.

### 3.2 Knee joint moments and stiffness

The knee joint ROM, moments, and stiffness are summarized in **Table 3**. The results indicate that external load conditions and joint angles exert significant and distinct effects on the biomechanical characteristics of the knee. In the sagittal plane, a highly significant main effect was observed for the angle (F = 75.536, P < 0.001, *η*^2^ = 0.744), whereas the main effect of external load and the interaction were not significant. In the coronal plane, a significant interaction existed between external load and angle (F = 13.113, P = 0.001, *η*^2^ = 0.335). Simple effect analysis revealed significant differences between external load conditions specifically at the 90° angle (P = 0.002). In the horizontal plane, significant main effects were found for both external load (F = 5.068, P = 0.033, *η*^2^ = 0.163) and angle (F = 23.445, P < 0.001, *η*^2^ = 0.474), though their interaction was not significant. Joint moments exhibited distinct patterns across the three anatomical axes. In the sagittal plane, a highly significant main effect of angle was observed (F = 53.349, P < 0.001, *η*^2^ = 0.672), with moments at 60° being significantly higher than at 90°. In the coronal plane, peak positive moments were not significantly affected by either external load or angle, suggesting that the tested conditions had limited effects on coronal-plane peak moments. In the horizontal plane, both external load (F = 5.527, P = 0.027, *η*^2^ = 0.175) and angle (F = 71.182, P < 0.001, *η*^2^ = 0.732) showed significant main effects, with moments decreasing as the joint angle increased. Regarding joint stiffness, a significant main effect of external load was observed in the sagittal plane (F = 8.864, P = 0.006, *η*^2^ = 0.254), indicating that the 10kg load significantly increased anteroposterior stiffness. In the coronal plane, a significant interaction between external load and angle was found (F = 9.252, P = 0.005, *η*^2^ = 0.262); simple effect analysis showed that external load at 90° led to a significant decrease in mediolateral stiffness (P = 0.012). In the horizontal plane, a significant interaction was observed (F = 32.906, P < 0.001, *η*^2^ = 0.559), where the 98 N external load resulted in a significant reduction in rotational stiffness across all angles.

**Table 3.**
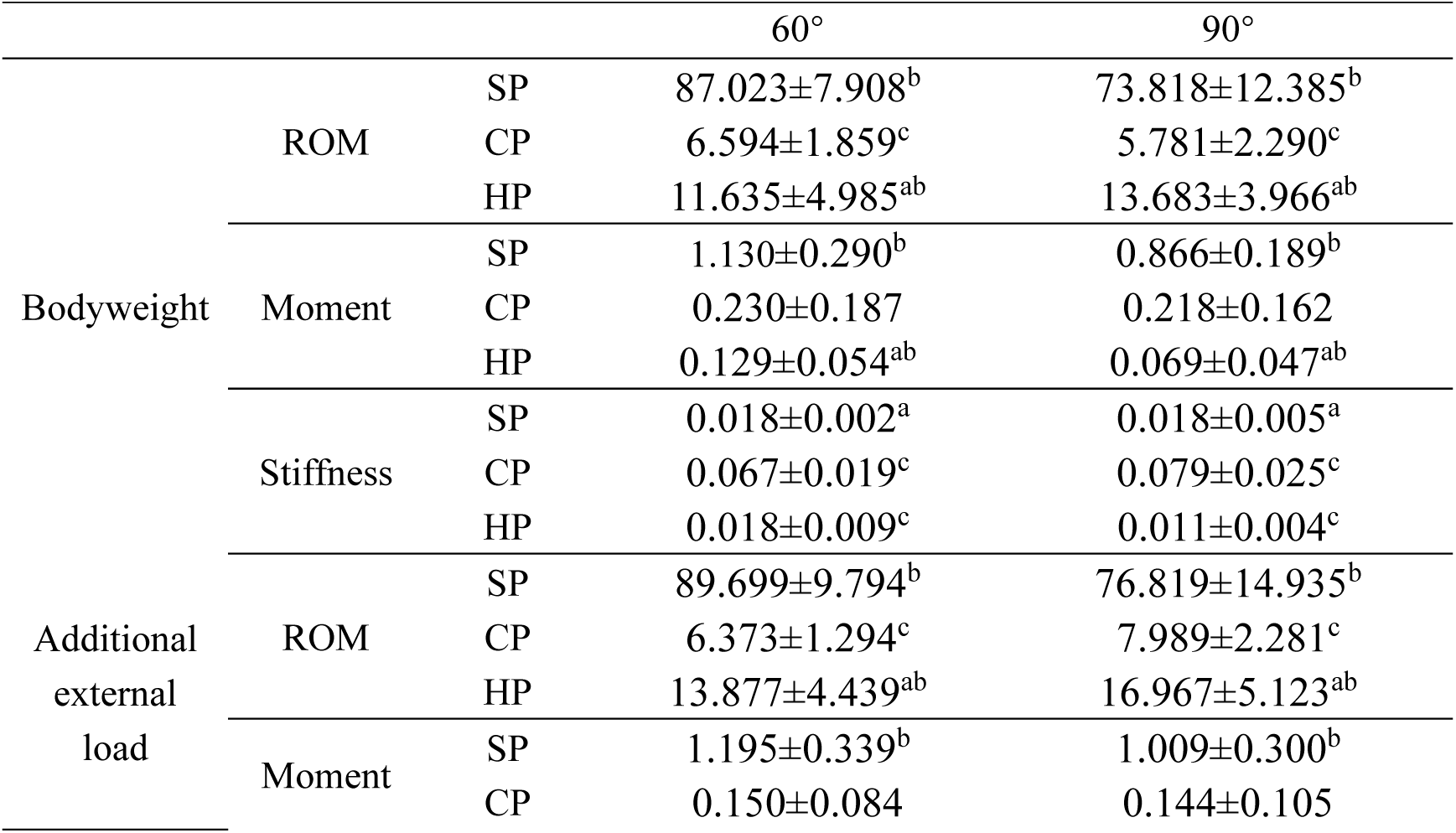
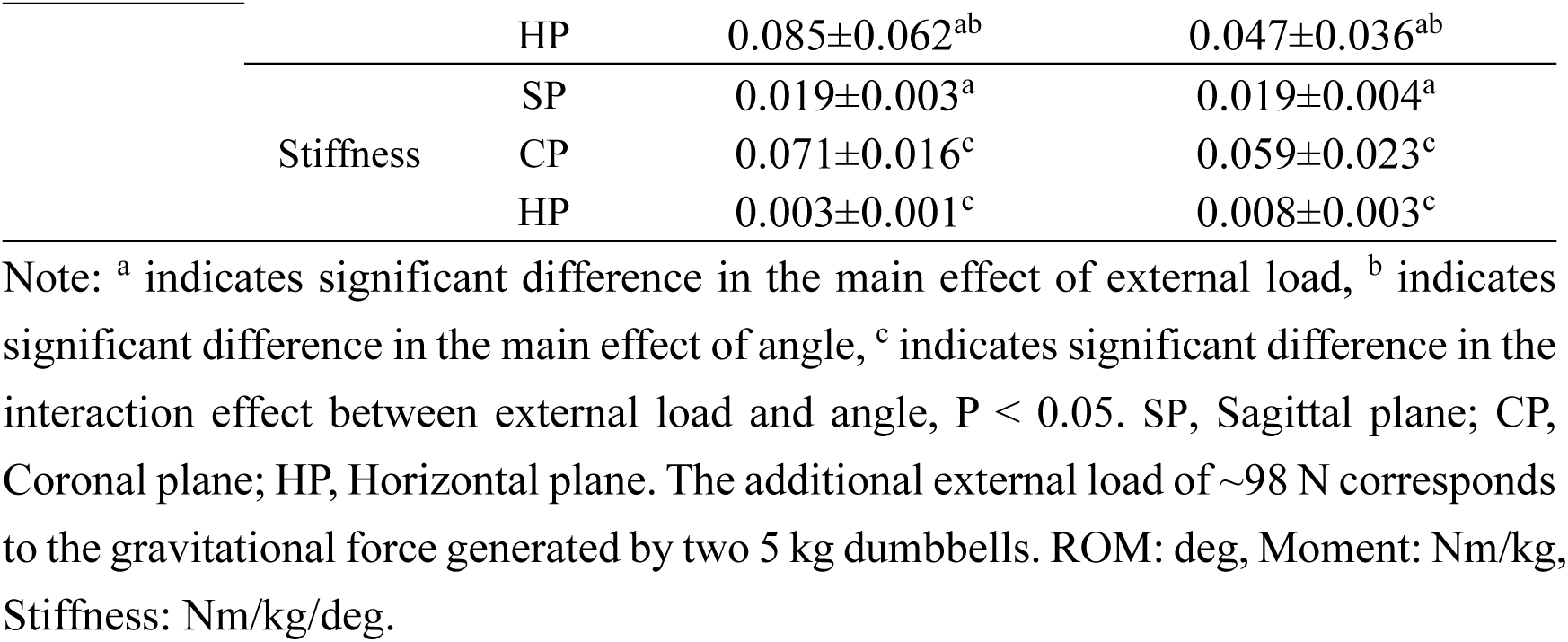
Joint ROM, peak joint moments and stiffnesses during Sagittal, coronal and Horizontal plane.

### 3.3 FEA of joint displacement and stress distribution in the knee joint

**Figure 3** shows the displacement distribution and equivalent (von-Mises) stress distribution of the knee joint finite element model under the bodyweight condition at 90° of flexion. The total displacement of the knee joint peaked at 3.01 mm, with a recorded range across the model of 0.01–3.01 mm. Deformation was highly concentrated at the posterior horn of the menisci, accurately replicating the posterior compression mechanism of the femoral condyles against the menisci during high-flexion postures. Stress analysis showed that the femoral cartilage had a maximum stress of 10.05 MPa, which showed up as clear regions of localized stress concentration. The internal equivalent stress in the menisci was between 0.00 and 16.82 MPa, with the highest stress being around five times higher than that of the tibial cartilage (3.36 MPa). The menisci have a lot more stress than the femoral and tibial cartilage. This means that they are good at spreading axial stresses through their circumferential hoop stiffness.

**Figure 3.**
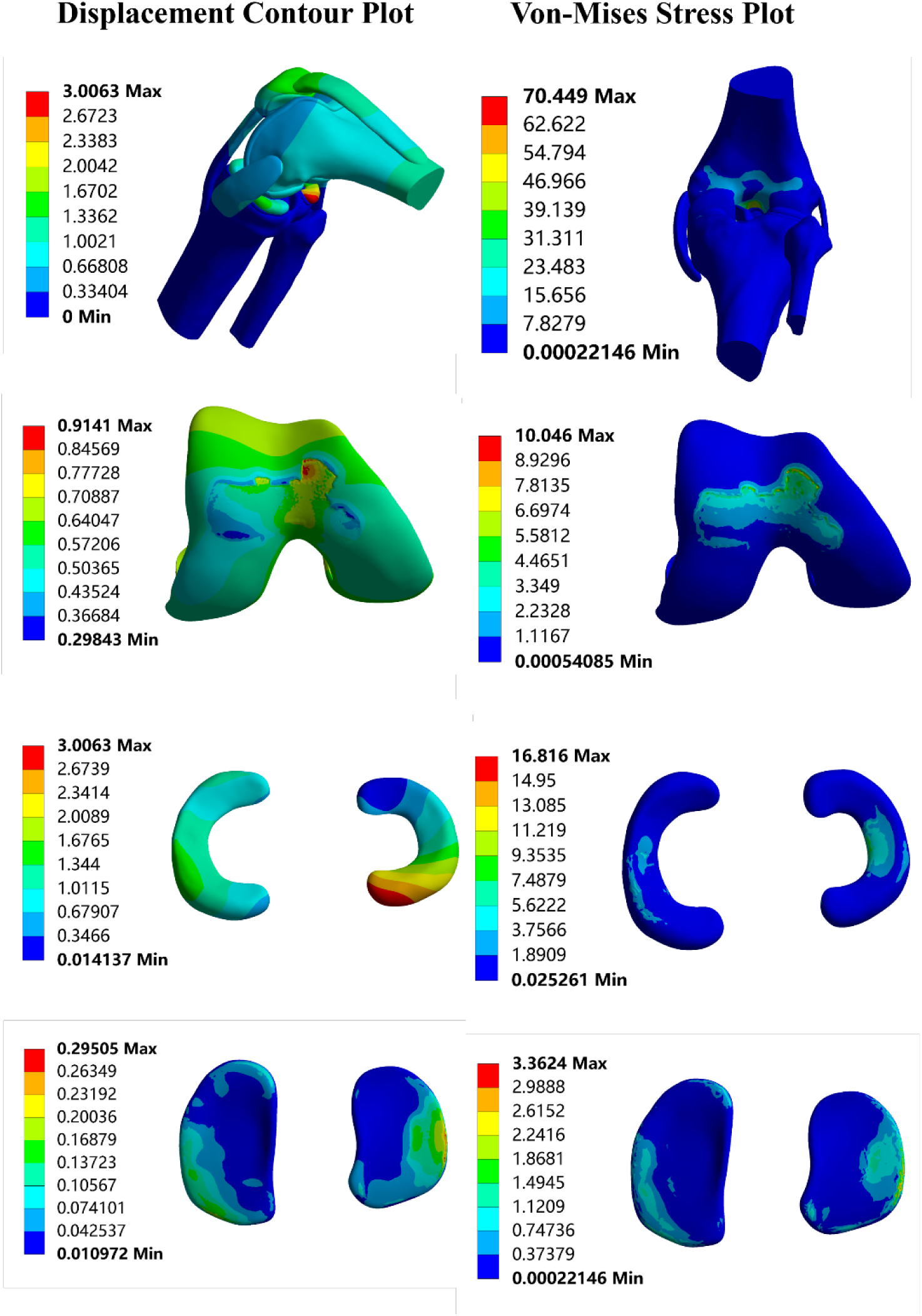
Displacement distribution plot and equivalent (von-Mises) stress distribution plot for the overall knee joint, femoral cartilage, tibial cartilage, and meniscus under the self-weight condition.

Under the high-intensity condition of a 98 N loaded lunge at 90° of flexion in **Figure 4**, the applied quadriceps force was 2490 N with a gravitational load of 504 N. The total displacement vector of the knee joint ranged from 0.01 to 3.12 mm. Under the 98 N external-load condition, the peak displacement (3.12 mm) was concentrated at the posterior horn of the menisci. The overall maximum equivalent stress reached 82.06 MPa, with the peak stress in the femoral cartilage reaching 11.62 MPa, showing a significant concentration toward the central contact area. In contrast, the peak stress in the tibial cartilage remained at a relatively low level of 3.49 MPa. The internal structures of the menisci experienced intense stress states, with the peak equivalent stress reaching 17.44 MPa, primarily distributed along the peripheral edges of the contact surface.

**Figure 4.**
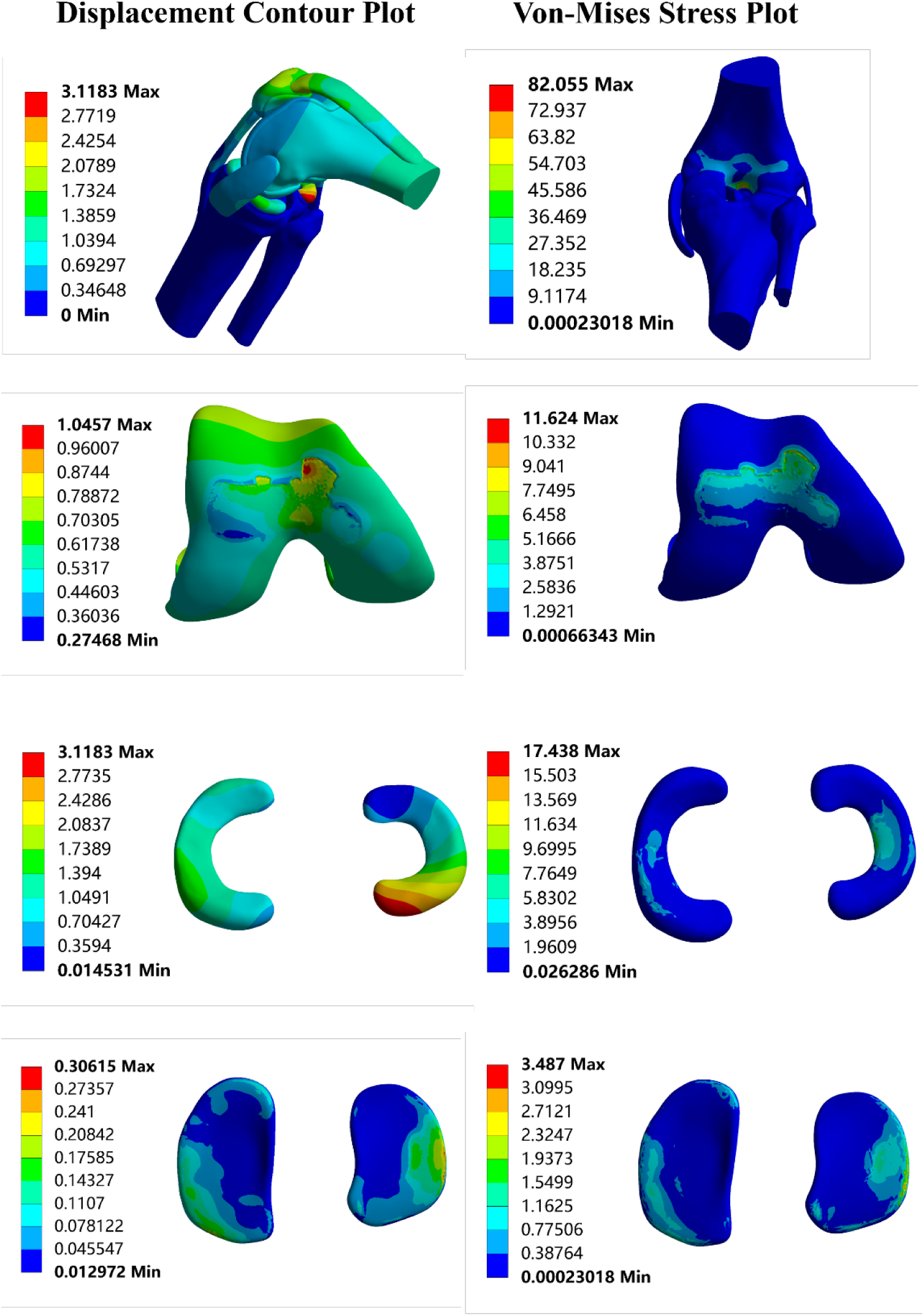
Displacement distribution plot and equivalent (von-Mises) stress distribution plot for the overall knee joint, femoral cartilage, tibial cartilage, and meniscus under the 10kg load condition

## 4. Discussion

This study relied on musculoskeletal modeling and FEA to investigate at how the knee joint works biomechanically during lunge exercises with different external load conditions and flexion angles. The results show the ways joint moments, stress distribution, and joint stiffness change depending on the type of motion. This provides valuable data to support sports training and rehabilitation programs.

This study demonstrates that variations in external load and flexion angles exert a significant influence on the biomechanical response of the knee. The external load altered knee joint mechanical demands, as reflected by changes in ROM and joint stiffness. Together with the angle-related changes in knee moments, these findings support the first hypothesis that tibial angle and external loading influence knee joint moments and stiffness. The tibial angle serves as the core factor determining the dynamic external load in the sagittal plane. Our data indicate that as the tibial angle decreased from 90° to 60°, the ROM in the sagittal and coronal planes increased substantially, accompanied by a highly significant rise in sagittal moments. Specifically, at a 60° flexion angle, the knee joint moments increased significantly, reaching peak values in the sagittal plane under the 10 kg external load condition; a notable increase was also observed in the horizontal plane. This phenomenon can be explained by the lever principle: as the anterior tilt of the tibia increases, the horizontal moment arm of the ground reaction force vector relative to the knee joint center increases significantly. To maintain postural balance, the knee extensors must generate greater contractile force to counteract the drastically increased gravitational moment. This finding aligns with the classic study by Escamilla et al. (Escamilla et al. 2008), which concluded that deeper lunges impose higher functional demands on the extensor mechanism. Other research similarly notes that as the angle between the lead tibia and the ground decreases, the lead knee ROM and hip flexion moments increase; if hip strength is sufficient, it may compensatorily reduce knee pressure (Schütz et al. 2012). These findings may have potential implications for load management during lunge-based training and rehabilitation, and tibial angle control may be considered as a strategy to regulate sagittal-plane knee mechanical demand.

Adding a 98 N external load significantly increased knee ROM in both the horizontal and coronal planes. This was accompanied by a dramatic drop in rotational stiffness. At various flexion angles, external load not only augmented joint moments but also markedly influenced joint stiffness. A decrease in joint stiffness indicates reduced resistance to angular displacement. Therefore, the effect of external load was mainly reflected in altered stiffness in the coronal and horizontal planes. In the coronal plane specifically, the statistical analysis revealed a significant interaction between load and angle, with the 98 N external load leading to a marked decrease in mediolateral stiffness at 90° flexion. This reduction in stiffness was accompanied by the FE finding that the posterior horn of the meniscus showed a peak displacement of 3.12 mm under the same condition. Together, these findings suggest that external loading may increase mechanical demand on the posterior meniscus during deep lunges, although this should not be interpreted as direct evidence of meniscal injury. Therefore, high-load lunge training should be prescribed cautiously, particularly when the goal is to control knee joint loading during rehabilitation or sports training.

In regard to Hypothesis 2, this study further examined the influence of external load on internal stress and displacement within knee joint tissues. Under the 90° tibial-angle condition with a 98 N external load, the FE results showed posterior concentration of displacement and stress in the meniscus. This flexion-related posterior meniscal loading pattern is consistent with previous FE findings on tibiofemoral mechanics and meniscal stress distribution under high-flexion conditions (Kedgley et al. 2019; Zhang et al. 2021). The peak displacement occurred in the posterior horn of the meniscus, reaching 3.12 mm, while the peak equivalent stress in the meniscus reached 17.44 MPa, nearly five times that of the tibial cartilage. The maximum stress in the femoral cartilage was 11.62 MPa, whereas the tibial cartilage stress remained relatively modest at 3.49 MPa. These findings are consistent with the second hypothesis that external loading increases localized tissue-level mechanical responses under the 90° condition. However, they should be interpreted as potential biomechanical indicators of mechanically unfavorable loading rather than direct evidence of meniscal injury.

While linear elastic models ensure computational efficiency, they may not fully capture the complex behaviors of cartilage and menisci under dynamic loading. As highlighted in previous studies (Galbusera et al. 2014; Wang et al. 2015), these models may overestimate peak stresses compared to more advanced models. Future studies will integrate more sophisticated material models.

The combined musculoskeletal modeling and FEA framework provides a multiscale approach for linking joint-level mechanical outputs with tissue-level stress and displacement patterns. By integrating experimentally driven joint loading with tissue-level FE responses, this framework offers useful biomechanical evidence for understanding how tibial angle and external load influence knee tissue loading during lunges. Therefore, the findings may help inform load management and exercise prescription in rehabilitation and sports training, while the tissue-level results should still be interpreted in the context of the representative-subject FE model and simplified material assumptions.

## 5. Limitation

Although this study provides important insights into the biomechanical characteristics of the knee joint under different loads and flexion angles, several limitations remain. Firstly, the FE model was developed from CT and MRI data from one representative participant; therefore, it does not account for individual variability, such as sex-specific differences, athletic ability, and clinical pathological states. Secondly, large soft tissue deformations in deep flexion made numerical convergence difficult, preventing this study from performing a quantitative micro-mechanical comparison at a 60° tibial angle. Furthermore, this study focuses on simulating knee joint biomechanics using musculoskeletal modeling and FEA, which may not fully capture the dynamic complexities of knee motion. The material properties for certain knee components were assumed based on typical values from the literature, and the model does not account for variations in these properties across individuals. Moreover, the ligaments and menisci were modeled as isotropic linear elastic materials, although they are fiber-reinforced and anisotropic in vivo. Therefore, the FE results should be interpreted mainly as relative stress and displacement patterns rather than exact tissue-level responses. Finally, the study only evaluated fixed loads and specific knee flexion degrees, not a wider range of motion situations. Large soft-tissue deformation and mesh distortion may also affect computational precision. Future research should include multi-sample clinical cohorts, more complex material models, and dynamic simulation improvements to enhance the generalizability and clinical applicability of the conclusions.

## 6. Conclusion

This study integrated musculoskeletal modeling with finite element analysis to evaluate knee joint biomechanics during forward lunges performed with different tibial angles and external load conditions. The results indicate that tibial angle is the primary factor affecting sagittal-plane knee loading, while external load mainly impacts joint stiffness in the coronal and horizontal planes. The addition of a 98 N external load was associated with diminished rotational and lateral stiffness, indicating a lower capacity to resist angular displacement in the coronal and horizontal planes. Under the 90° loaded condition, stress and displacement became concentrated in the posterior horn of the meniscus. These findings indicate that deep, loaded lunges may create a mechanically unfavorable loading pattern for the posterior meniscus and should therefore be considered cautiously in rehabilitation and sports training.

## Author Contributions

All the authors contributed substantially to the manuscript. L.G., S.G., Z.L., and Z.G. were responsible for the conceptualization. L.G., S.G., Z.L., and Z.G. were responsible for investigation and methodology. L.G., G. F., and Z.G. were responsible for formal analysis and writing – original draft. S.G., L.G. and G. F. were responsible for writing – review & editing and supervision. All authors have read, provided feedback, and approved the submitted version.

## Ethical approval

This study has received full approval from the NBDX Ethics Committee (TY2025087) and strictly adhered to the informed consent process.

## References

Al Mohammad B, Gharaibeh MA. 2024. Magnetic resonance imaging of anterior cruciate ligament injury. Orthopedic Research and Reviews.233–242. doi:10.2147/ORR.S450336.

Alkjær T et al. 2020. Forward lunge before and after anterior cruciate ligament reconstruction: Faster movement but unchanged knee joint biomechanics. Plos one. 15(1):e0228071. doi:10.1371/journal.pone.0228071.

Andriacchi TP et al. 2004. A framework for the in vivo pathomechanics of osteoarthritis at the knee. Annals of biomedical engineering. 32(3):447–457.

Arias JT, Ramos J, Gallego A, Martinez-Cano JP. 2025. Incidence of injuries in the colombian medical football championship 2022: A prospective observational cohort study. Physical Activity and Health. 9(1). doi:10.5334/paah.502.

Beidokhti HN et al. 2017. The influence of ligament modelling strategies on the predictive capability of finite element models of the human knee joint. Journal of biomechanics. 65:1–11. doi:10.1016/j.jbiomech.2017.08.030.

Benos L, Stanev D, Spyrou L, Moustakas K, Tsaopoulos DE. 2020. A review on finite element modeling and simulation of the anterior cruciate ligament reconstruction. Frontiers in Bioengineering and Biotechnology. 8:967.

Chuter VH, de Jonge XAJ. 2012. Proximal and distal contributions to lower extremity injury: A review of the literature. Gait & posture. 36(1):7–15. doi:10.1016/j.gaitpost.2012.02.001.

Escamilla RF et al. 2008. Patellofemoral joint force and stress between a short-and long-step forward lunge. Journal of Orthopaedic & Sports Physical Therapy. 38(11):681–690. doi:10.2519/jospt.2008.2694.

Faul F, Erdfelder E, Lang A-G, Buchner A. 2007. G* power 3: A flexible statistical power analysis program for the social, behavioral, and biomedical sciences. Behavior research methods. 39(2):175–191.

Felson DT. 2013. Osteoarthritis as a disease of mechanics. Osteoarthritis and cartilage. 21(1):10–15. doi:10.1016/j.joca.2012.09.012.

Galbusera F et al. 2014. Material models and properties in the finite element analysis of knee ligaments: A literature review. Frontiers in bioengineering and biotechnology. 2:54. doi:10.3389/fbioe.2014.00054.

Gao L, Lu Z, Liang M, Baker JS, Gu Y. 2022. Influence of different load conditions on lower extremity biomechanics during the lunge squat in novice men. Bioengineering. 9(7):272. doi:10.3390/bioengineering9070272.

Gao L et al. 2023. Biomechanical effects of exercise fatigue on the lower limbs of men during the forward lunge. Frontiers in Physiology. 14. doi:10.3389/fphys.2023.1182833.

Gao Z et al. 2025. Musculoskeletal modelling sequentially integrated with stress simulation reveals asymmetrical knee loading and ligament stress during long-distance running. BMC Sports Science, Medicine and Rehabilitation. 17(1):337. doi:10.1186/s13102-025-01372-3.

Haut Donahue TL, Hull M, Rashid MM, Jacobs CR. 2002. A finite element model of the human knee joint for the study of tibio-femoral contact. J Biomech Eng. 124(3):273–280. doi:10.1115/1.1470171.

Hermens HJ, Freriks B, Disselhorst-Klug C, Rau G. 2000. Development of recommendations for semg sensors and sensor placement procedures. Journal of electromyography and Kinesiology. 10(5):361–374. doi:10.1016/S1050-6411(00)00027-4.

Hsue B-J, Miller F, Su F-C. 2009. The dynamic balance of the children with cerebral palsy and typical developing during gait. Part i: Spatial relationship between com and cop trajectories. Gait & posture. 29(3):465–470. doi:10.1016/j.gaitpost.2008.11.007.

Innocenti B, Salandra P, Pascale W, Pianigiani S. 2016. How accurate and reproducible are the identification of cruciate and collateral ligament insertions using mri? The Knee. 23(4):575–581. doi:10.1016/j.knee.2015.07.015.

Kedgley AE et al. 2019. Predicting meniscal tear stability across knee-joint flexion using finite-element analysis. Knee Surgery, Sports Traumatology, Arthroscopy. 27(1):206–214. doi:10.1007/s00167-018-5090-4.

Kiapour A et al. 2014a. Finite element model of the knee for investigation of injury mechanisms: Development and validation. 136(1):011002.

Kiapour AM et al. 2014b. Diagnostic value of knee arthrometry in the prediction of anterior cruciate ligament strain during landing. The American journal of sports medicine. 42(2):312–319. doi:10.1177/0363546513509961.

Li F et al. 2025. Dynamic simulation of knee joint mechanics: Individualized multi-moment finite element modelling of patellar tendon stress during landing. Journal of Biomechanics. 186:112730. doi:10.1016/j.jbiomech.2025.112730.

Li G, Gil J, Kanamori A, Woo S-Y. 1999. A validated three-dimensional computational model of a human knee joint. doi:10.1115/1.2800871.

Lu Z et al. 2025. Computationally tuned dual-layer lattice pads adapted to gait-induced pressure distribution. npj Advanced Manufacturing. 2(1):43. doi:10.1038/s44334-025-00055-8.

Lu Z, Sun D, Kovács B, Radák Z, Gu Y. 2023. Case study: The influence of achilles tendon rupture on knee joint stress during counter-movement jump–combining musculoskeletal modeling and finite element analysis. Heliyon. 9(8). doi:10.1016/j.heliyon.2023.e18410.

Mei Q et al. 2022. Dataset of lower extremity joint angles, moments and forces in distance running. Heliyon. 8(11):e11517. doi:10.1016/j.heliyon.2022.e11517.

Mei Q, Gu Y, Xiang L, Baker JS, Fernandez J. 2019. Foot pronation contributes to altered lower extremity loading after long distance running. Frontiers in Physiology. 10:573. doi:10.3389/fphys.2019.00573.

Nakagawa S, Cuthill IC. 2007. Effect size, confidence interval and statistical significance: A practical guide for biologists. Biological reviews. 82(4):591–605. doi:10.1111/j.1469-185X.2007.00027.x.

Navacchia A et al. 2018. Loading and kinematic profiles for patellofemoral durability testing. Journal of the mechanical behavior of biomedical materials. 86:305–313. doi:10.1016/j.jmbbm.2018.06.035.

Pena E, Calvo B, Martinez M, Doblare M. 2006. A three-dimensional finite element analysis of the combined behavior of ligaments and menisci in the healthy human knee joint. Journal of biomechanics. 39(9):1686–1701. doi:10.1016/j.jbiomech.2005.04.030.

Rajagopal A et al. 2016. Full-body musculoskeletal model for muscle-driven simulation of human gait. IEEE transactions on biomedical engineering. 63(10):2068–2079. doi:10.1109/TBME.2016.2586891.

Rao H, Bakker R, McLachlin S, Chandrashekar N. 2024. Computational study of extrinsic factors affecting acl strain during single-leg jump landing. BMC Musculoskeletal Disorders. 25(1):318. doi:10.1111/os.13980.

Ren S et al. 2022. Finite element analysis and experimental validation of the anterior cruciate ligament and implications for the injury mechanism. Bioengineering. 9(10):590. doi:10.3390/bioengineering9100590.

Riemann BL, Lapinski S, Smith L, Davies G. 2012. Biomechanical analysis of the anterior lunge during 4 external-load conditions. Journal of athletic training. 47(4):372–378. doi:10.4085/1062-6050-47.4.16.

Rooks NB, Besier TF, Schneider MT. 2022. A parameter sensitivity analysis on multiple finite element knee joint models. Frontiers in bioengineering and biotechnology. 10:841882. doi:10.3389/fbioe.2022.841882.

Schütz P, List R, Zemp R, Lorenzetti S. Influence of the step length and position of the front knee on the load conditions of the knee and hip during lunges. In: ISBS-Conference Proceedings Archive. p 406–409.

Shao B, Xing J, Zhao B, Wang T, Mu W. 2022. Role of the proximal tibiofibular joint on the biomechanics of the knee joint: A three-dimensional finite element analysis. Injury. 53(7):2446–2453. doi:10.1016/j.injury.2022.05.027.

Smith BE et al. 2018. Incidence and prevalence of patellofemoral pain: A systematic review and meta-analysis. PloS one. 13(1):e0190892. doi:10.1371/journal.pone.0190892.

van Leeuwen GJ, de Schepper EI, Bindels PJ, Bierma-Zeinstra SM, van Middelkoop M. 2023. Patellofemoral pain in general practice: The incidence and management. Family Practice. 40(4):589–595. doi:10.1093/fampra/cmad087.

Wang Y, Li Z, Wong DW-C, Zhang M. 2015. Effects of ankle arthrodesis on biomechanical performance of the entire foot. PloS one. 10(7):e0134340. doi:10.1371/journal.pone.0134340.

Ward K, Kavanagh R, O’Connor S, Cooper D. 2026. ‘Beginner to 5kg’: A 6-week novel hybrid (online and in-person) strength training and health education programme for middle-aged and older women. Physical Activity and Health. 10(1).

Wu HW, Tsai CF, Liang KH, Chang YW. 2020. Effect of loading devices on muscle activation in squat and lunge. J Sport Rehabil. 29(2):200–205. doi:10.1123/jsr.2018-0182.

Yagi M et al. 2002. Biomechanical analysis of an anatomic anterior cruciate ligament reconstruction. The American journal of sports medicine. 30(5):660–666. doi:10.1177/03635465020300050501.

Yan M et al. 2024. Model properties and clinical application in the finite element analysis of knee joint: A review. 16(2):289–302. doi:10.1111/os.13980.

Yang S et al. 2024. Stress and strain changes of the anterior cruciate ligament at different knee flexion angles: A three-dimensional finite element study. Journal of Orthopaedic Science. 29(4):995–1002. doi:10.1016/j.jos.2023.05.015.

Yu L et al. 2023. Dose–response effect of incremental lateral-wedge hardness on the lower limb biomechanics during typical badminton footwork. Journal of Sports Sciences. 41(10):972–989. doi:10.1080/02640414.2023.2257513.

Zhang X, Yuan S, Wang J, Liao B, Liang D. 2021. Biomechanical characteristics of tibio-femoral joint after partial medial meniscectomy in different flexion angles: A finite element analysis. BMC Musculoskeletal Disorders. 22(1):322. doi:10.1186/s12891-021-04187-8.

Zhou H et al. A pilot study of muscle force between normal shoes and bionic shoes during men walking and running stance phase using opensim. In: Actuators. Multidisciplinary Digital Publishing Institute. Vol. 10 p 274.

